# scite: a smart citation index that displays the context of citations and classifies their intent using deep learning

**DOI:** 10.1101/2021.03.15.435418

**Authors:** J.M. Nicholson, M. Mordaunt, P. Lopez, A. Uppala, D. Rosati, N.P. Rodrigues, P. Grabitz, S.C. Rife

**Affiliations:** scite, Brooklyn, NY, USA; science-miner, France; Charite Universitaetsmedizin Berlin, Berlin, Germany; Murray State University, Murray, KY, USA

**Author notes:** Address correspondence to: Joshua M. Nicholson, PhD, scite Inc., 334 Leonard St, #6, Brooklyn, NY 11211, USA.

## Abstract

Citation indices are tools used by the academic community for research and research evaluation which aggregate scientific literature output and measure scientific impact by collating citation counts. Citation indices help measure the interconnections between scientific papers but fall short because they only display paper titles, authors, and the date of publications, and fail to communicate contextual information about why a citation was made. The usage of citations in research evaluation without due consideration to context can be problematic, if only because a citation that disputes a paper is treated the same as a citation that supports it. To solve this problem, we have used machine learning and other techniques to develop a “smart citation index” called scite, which categorizes citations based on context. Scite shows how a citation was used by displaying the surrounding textual context from the citing paper, and a classification from our deep learning model that indicates whether the statement provides supporting or disputing evidence for a referenced work, or simply mentions it. Scite has been developed by analyzing over 23 million full-text scientific articles and currently has a database of more than 800 million classified citation statements. Here we describe how scite works and how it can be used to further research and research evaluation.

## Introduction

Citations are a critical component of scientific publishing, linking research findings across time. The first citation index in science, created in 1960 by Eugene Garfield and the Institute for Scientific Information, aimed to “be a spur to many new scientific discoveries in the service of mankind” (*1*). Citation indices have facilitated the discovery and evaluation of scientific findings across all fields of research. Citation indices have also led to the establishment of new research fields such as bibliometrics, scientometrics, and quantitative studies, which have been informative in better understanding science as an enterprise. From these fields have come a variety of citation-based metrics like the H-index, a measurement of researcher impact (*2*), the Journal Impact Factor (JIF), a measurement of journal impact (*3, 4*), and the citation count, a measurement of article impact. Despite the widespread use of bibliometrics, there has been few improvements in citations and citation indices themselves. Such stagnation might be due partly to the fact that citations and publications are largely behind paywalls, thus creating a citation index, let alone to introducing innovations in citations or citation indices, is exceedingly difficult and can be prohibitively expensive. This trend is changing, however, with open access publications becoming the standard (*5*) and organizations such as the Initiative for Open Citations (*6*) helping to make citations open. Additionally, with millions of documents being published each year, creating a citation index is a large-scale challenge involving significant financial and computational costs.

Historically, citation indices have only shown the connections between scientific papers without any further contextual information such as why a citation was made. Because of the lack of context and limited metadata available beyond paper titles, authors, and the date of publications it has only been able to calculate how many times a work has been cited, not analyze broadly *how* it has been cited. This is problematic given citations’ central role in the evaluation of research. In short, not all citations are made equally, yet we’ve been limited to treating them as such.

Here we describe scite (scite.ai), a new citation index and tool that takes advantage of recent advances in artificial intelligence to produce “Smart Citations.” Smart Citations reveal how a scientific paper has been cited by providing the context of the citation and a classification system describing whether it provides supporting or disputing evidence for the cited claim or if it just mentions it.

Such enriched citation information is more informative than a traditional citation index. For example, when Vigano et. al (*7*) cites Nicholson et. al (*8*), traditional citation indices report this citation by displaying the title of the citing paper and other bibliographic information such as the journal, year published, and other metadata (Figure 1). Traditional citation indices do not have the capacity to examine contextual information or how the citing paper used the citation, such as whether it was made to support or dispute the findings of the cited paper or if it was made in the introduction or the discussion section. Smart Citations display the same bibliographical information displayed in traditional citation indices and provide contextual information such as the citation statement (the sentence containing the in-text citation from the citing article), the citation context (the sentence before and after the citation statement), the location of the citation within the citing article (Introduction, Materials and Methods, Results, Discussion, etc.), the citation type indicating intent (supporting, disputing, or mentioning), and editorial information such as corrections and whether the article has been retracted (Figure 2).

**Figure 1.**
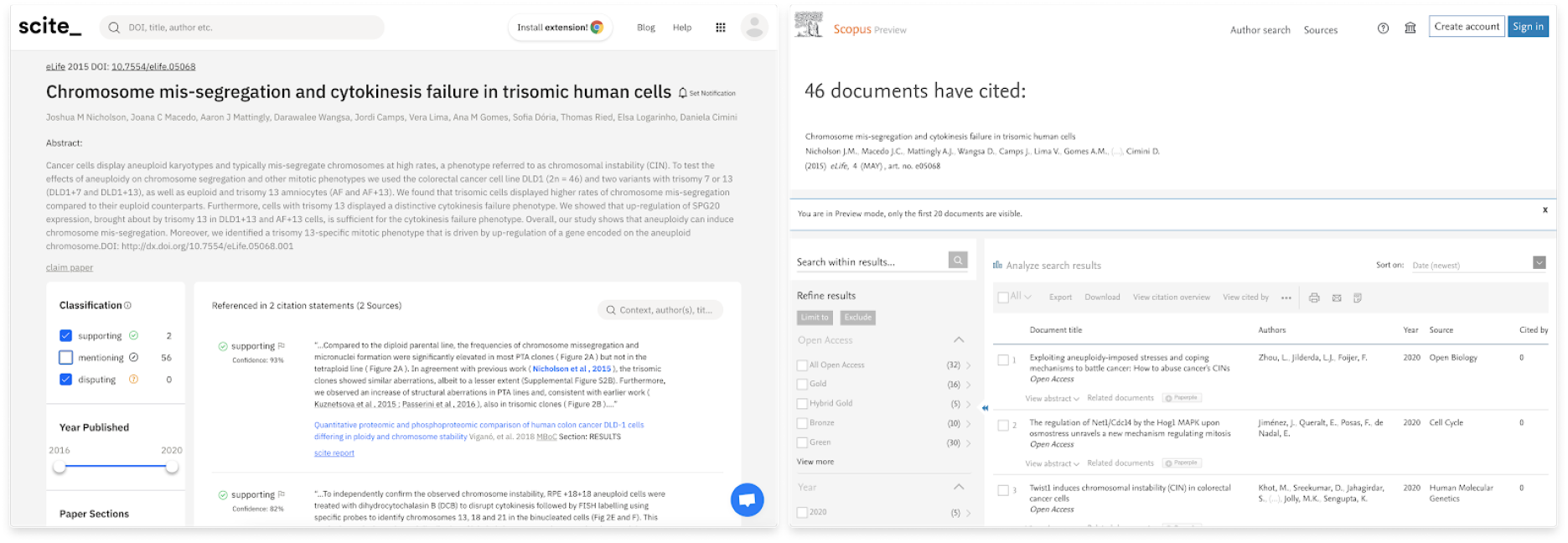
A smart citation report (left) versus a traditional citation report (right). The traditional citation report contains article metadata such as title, author(s), journal, date, and persistent identifier whereas the smart citation index displays the same information as well as the citation context, citation type, and citation location.

**Figure 2.**
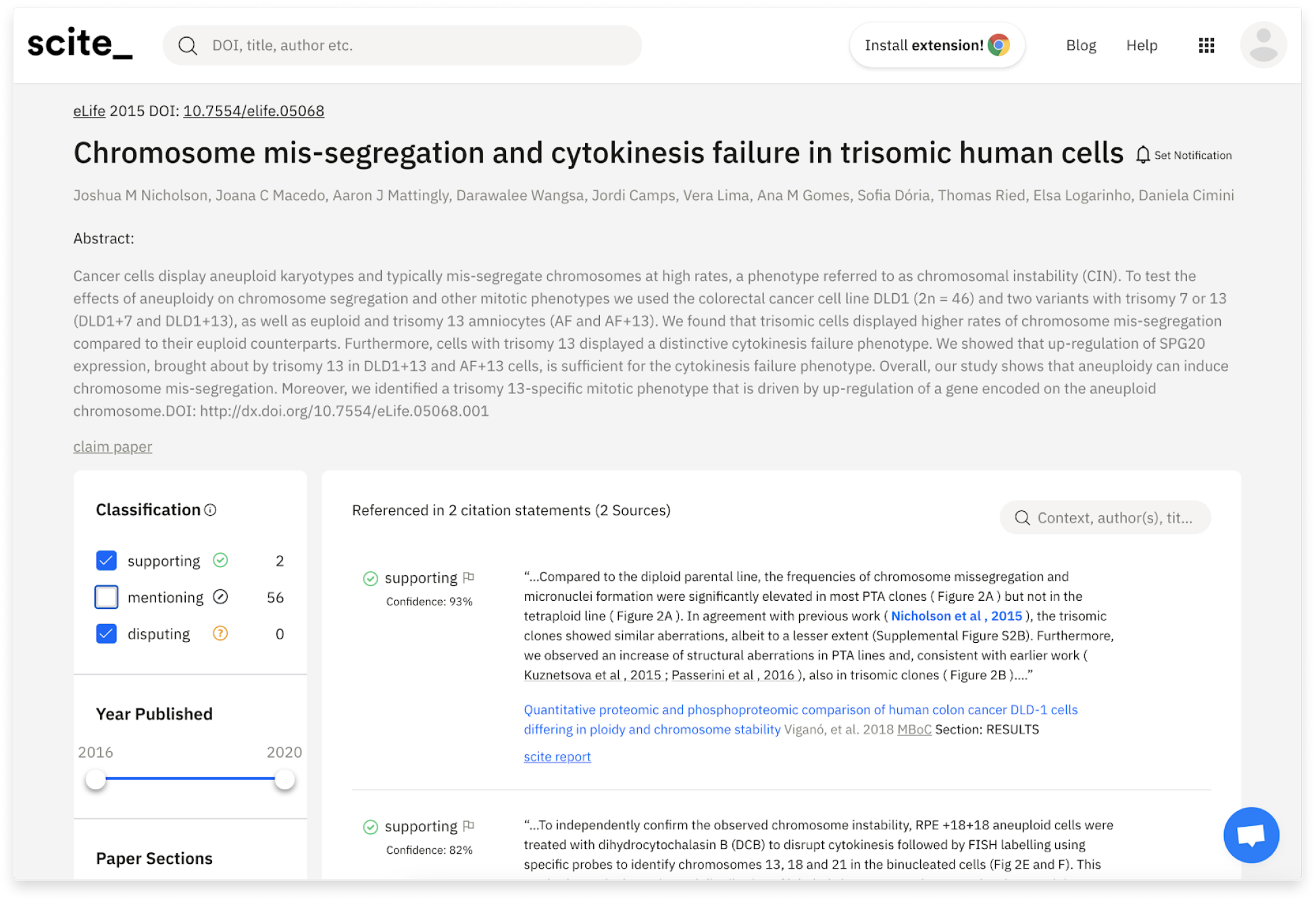
Example of scite report page. The scite report page shows citation context, citation type, and various features used to filter and organize this information.

Adding such information to citation indices has been proposed before. In 1959, Garfield described an “intelligent machine” to produce “citation markers,” such as “critique” or, jokingly, “calamity for mankind.” (*1*) Citation types describing various uses of citations have been systematically described by Peroni and Shotton in CiTO, the *C*itation *T*yping *O*ntology (*9*). Researchers have used these classifications or variations of them in several bibliometric studies, such as the analysis of citations (*10*) made to the retracted Wakefield paper (*11*), which found most citations to be negative in sentiment. Leung et. al (*12*) analyzed the citations made to a five-sentence letter purporting to show opioids as non-addictive (*13*), finding that most citations were uncritically citing the work. Based on these findings, the journal appended a warning to the original letter. In addition to citation analyses at the individual article level, citation analyses taking into account the citation type have also been performed on subsets of articles or even entire fields of research. Greenberg (*14*) discovered that citations were being distorted, e.g. used selectively to exclude contradictory studies to create a false authority in a field of research, a practice carried even into grant proposals. Selective citing might be malicious as suggested in the Greenberg study but it might also simply reflect sloppy citation practices or citing without reading. Indeed, Letrud and Hernes recently documented many cases where people were citing reports for the opposite conclusions the original authors made (*15*).

Despite the advantages of citation types, citation classification and analysis require substantial manual effort on the part of researchers to perform even small scale analyses (*16*). Automating the classification of citation types would allow researchers to dramatically expand the scale of citation analyses thereby allowing researchers to quickly assess large portions of scientific literature. PLOS Labs attempted to enhance citation analysis with the introduction of “rich citations,” which included various additional features to traditional citations such as retraction information and where the citation appeared in the citing paper (*17*). However, the project seemed to be mostly a proof of principle, and work on rich citations stopped in 2015. The Colil Database (*18*) and SciRide Finder (*19*) both allow users to see the citation context from open access articles indexed in Pubmed Central, yet adoption seems to be low on both tools, presumably due to limited coverage of only open access articles. In addition to the development of such tools to augment citation analysis, various researchers have performed automated citation typing; for example, Athar and Teufel used machine learning to classify and identify “negative citations” (*20*–*22*).

Here, by combining the largest citation type analysis performed to date and developing a useful user interface that takes advantage of the extra contextual information available, we introduce scite, a smart citation index.

## Method

### Overview

Smart citations are created by extracting and analyzing citation statements from full-text scientific articles. This process is broken into four major steps (see Figure 3): 1) the retrieval of scientific articles, 2) the identification and matching of in-text citations and references within a scientific article, 3) the matching of references against a bibliographic database, 4) the classification of the citation statements into citation types using deep learning. We describe the four components in more detail below.

**Figure 3.**
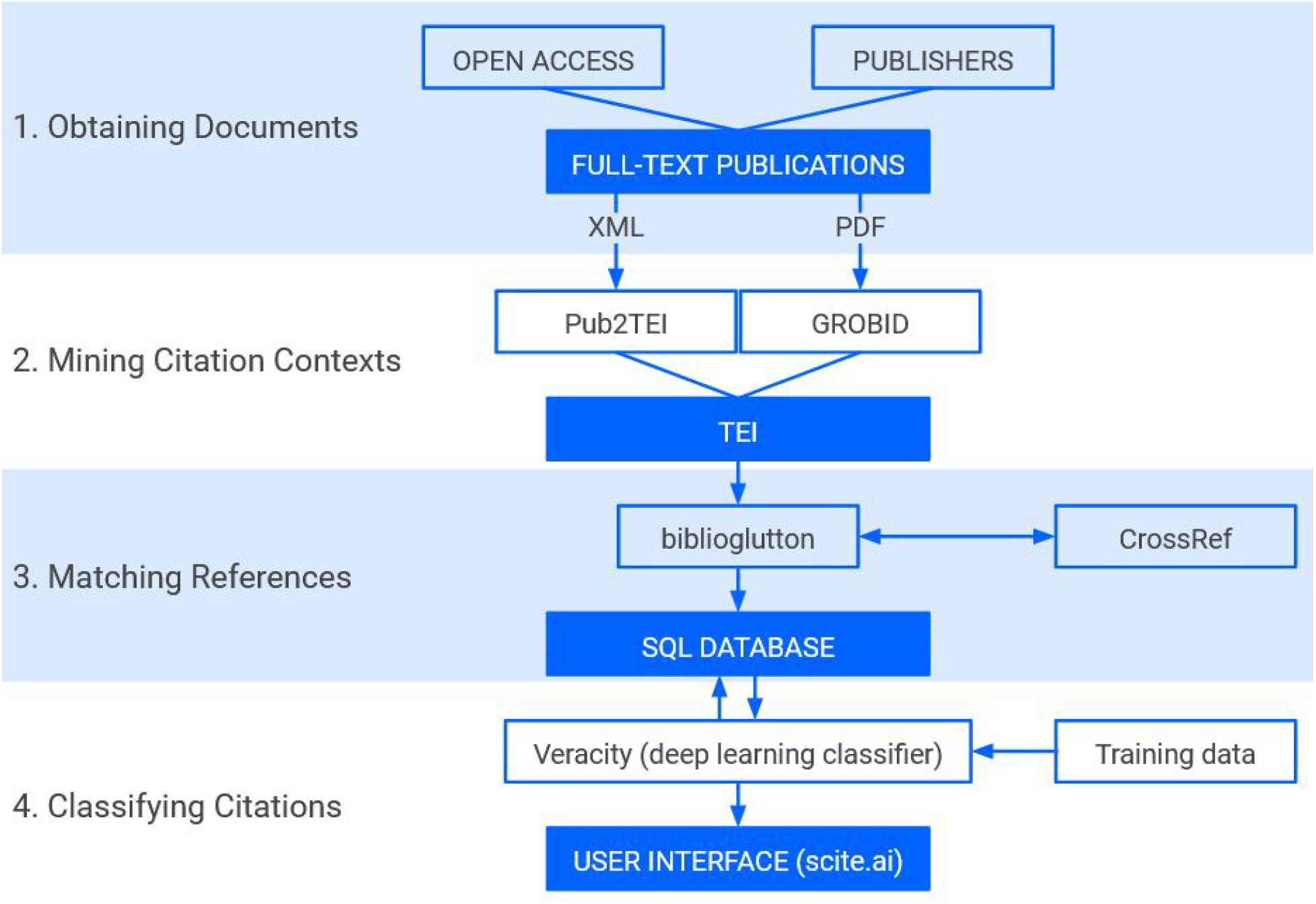
The scite ingestion process. Documents are retrieved from the Internet, as well as received through file transfers directly from publishers and other aggregators. They are then processed to identify citations, which are then tied to items in a paper’s reference list. Those citations are then verified, and the information is inserted into scite’s database.

### Retrieval of scientific documents

In order to extract and classify citation statements and the citation context, access to full-text scientific articles is necessary. We utilize open access repositories such as Pubmed Central and a variety of open sources as identified by Unpaywall (*23*) such as open access publishers websites, university repositories, and preprint repositories to analyze open access articles. Subscription articles used in our analyses have been made available through indexing agreements with over a dozen publishers including Wiley, BMJ, Karger, Sage, Europe PMC, Thieme, Cambridge University Press, Rockefeller University Press, IOP, Microbiology Society, Frontiers, and other smaller publishers. Once a source of publications is established documents are retrieved on a regular basis as new articles become available in order to keep the citation record fresh. Depending on the source, documents may be retrieved and processed anywhere between daily and monthly.

### Identification of in-text citations and references from PDF and XML documents

A large majority of scientific articles are only available as PDF files^1^, a format designed for visual layout and printing, not text-mining. In order to match and extract citation statements from PDFs with high fidelity, an automated process for converting PDF files into reliable structured content is required. Such conversion is challenging as it requires identifying in-text citations (the numerical or textual callouts that refer to a particular item in the reference list), identifying and parsing the full bibliographical references in the reference list, linking in-text citations to the correct items in this list, and linking these items to their digital object identifiers (DOIs) in a bibliographic database. Since our goal is to eventually process all scientific documents, this process must be scalable and affordable. To accomplish this, we utilize GROBID, an open-source PDF-to-XML converter tool for the scientific literature (*24*). The goal of GROBID is to automatically convert scholarly PDFs into structured XML representations suitable for large-scale analysis. The structuration process is realized by a cascade of supervised machine learning models. The tool is highly scalable (around 5 PDF documents per second on a 4-core server), robust, and includes a production-level web API, a Docker image, and benchmarking facilities. GROBID is used by many large scientific information service providers such as ResearchGate, CERN, and the Internet Archive to support their ingestion and document workflows (*25*). The tool is also used for creating machine-friendly datasets of research papers, for instance, the recent CORD-19 dataset (*26*).

Particularly relevant to scite, GROBID was benchmarked as the best Open Source bibliographical references parser by the recent study of (*27*) and has a relatively unique focus on citation context extraction at scale, as illustrated by its usage for building the large-scale S2ORC, a corpus of 380.5M citations including citation mentions excerpts from the full-text body (*28*).

In addition to PDFs, some scientific articles are available as XML files, for instance, the Journal Article Tag Suite (JATS) format. Formatting articles in PDF and XML has become standard practice for most mainstream publishers. While structured XML can solve many issues that need to be addressed with PDFs, XML full texts appear in a variety of different native publisher XML formats, often incomplete and inconsistent from one to another, loosely constrained, and evolving over time into specific versions.

To standardize the variety of XML formats we receive into a common format, we rely upon the open-source tool Pub2TEI (*29*). Pub2TEI converts various XML styles from publishers to the same standard TEI format as the one produced by GROBID. This centralizes our document processing across PDF and XML sources.

### Matching references against the bibliographic database Crossref

Once we have identified and matched the in-text citation to an item in a paper’s reference list, this information must be validated. We use an open-source tool, biblio-glutton (*30*), which takes a raw bibliographical reference, as well as optionally parsed fields (e.g., title, author names, etc.) and matches it against the Crossref database - widely regarded as the industry-standard source of ground truth for scholarly publications^2^. The matching accuracy of a raw citation reaches an F-score of 95.4 on a set of 17,015 raw references associated with a DOI, extracted from 1942 PMC articles (evaluation data and scripts are available on the project GitHub repository; see biblio-glutton, 2018-2020). In an end-to-end perspective, based on an estimate from PMC articles, combining GROBID PDF extraction of citations and bibliographical references with biblio-glutton validations, the pipeline successfully associates around 70% of citation contexts to cited papers with correctly identified DOIs in a given PDF file. When the full-text XML version of an article is available from a publisher, references and linked citation contexts are normally correctly encoded, and the proportion of fully solved citation contexts corresponding to the proportion of cited paper with correctly identified DOIs is around 95% for PMC XML JATS files.

### Task modeling and training data

Extracted citation statements are classified into supporting, disputing, or mentioning, in order to identify studies that have tested the claim and to evaluate how a scientific claim has been evaluated in the literature by subsequent research. We emphasize that scite is not doing sentiment analysis (*31, 22, 32, 33*), where a subjective polarity is associated with a claim, but a discrete classification into three discursive functions relative to the scientific debate (see Murray et al (*34*) for an example of previous work with typing citations based on rhetorical intention). We consider that for capturing the reliability of a claim, a classification decision into supporting or disputing must be evidence-based, backed by scientific arguments. For instance, a mere negative opinion (e.g., negative sentiment) about a cited work not supported by any scientific and/or technical facts, studies, or replication evidence will be classified as mentioning, while it would be classified as negative by sentiment analysis.

The main challenge of this classification task is the highly imbalanced distribution of the three classes. Based on manual annotations of different publication domains and sources, we estimate the average distribution of citation statements as 92.6% mentioning, 6.5% supporting, and 0.8% disputing statements. Obviously, the less frequent the class, the more valuable it is. Most of the efforts in the development of our automatic classification system have been all directed to address this imbalanced distribution. This task has required first the creation of original training data by experts– scientists with experience in reading and interpreting scholarly papers. Focusing on data quality, the expert classification was realized by multiple-blind manual annotation (at least two annotators working in parallel on the same citation), followed by a reconciliation step where the disagreements were further discussed and analyzed by the annotators. In order to keep track of the progress of our automatic classification over time, we created a holdout set of 9,708 classified citation records with a class distribution as close as possible to the actual distribution in current scholarly publications.

We developed separately a working set where we tried to oversample the two less frequent classes (supporting, disputing) with the objective of addressing the difficulties implied by the imbalanced automatic classification. We exploited the classification scores of our existing classifiers to select more likely supporting and disputed statements for manual classification. At the present time, this set contains 38,925 classified citation records. The automatic classification system was trained with this working set, and continuously evaluated with the immutable holdout set to avoid as much bias as possible. An n-fold cross-evaluation on the working set for instance would have been misleading because the distribution of the classes in this set was artificially modified to boost the classification accuracy of the less frequent classes.

Before reconciliation, the observed average Inter-Annotator-Agreement percentage was 78.5% in the open domain and close to 90% for batches in biomedicine. Reconciliation, further completed with expert review by core team members, resulted in highly consensual classification decisions, which contrast with typical multi-round disagreement rates observed with sentiment classification. Athar (*31*), for instance, reports Cohen’s *k* annotator agreement of 0.675 and Ciancarini et al. (*35*) reports *k*=0.13 and *k*=0.15 for the property groups covering *confirm/supports* and *critiques* citation classification labels. A custom doccano (*36*) web application (called *sciteano*) was deployed to support the first round of annotations.

Overall, the creation of our current training and evaluation holdout data sets has been a major two-year effort involving up to 8 expert annotators and nearly 50 thousand classified citation records. In addition to the class, each record includes the citation sentence, the full “snippet” (citation sentence plus previous and next sentences), the source and target DOI, the reference callout string, and the hierarchical list of section titles where the citation occurs.

### Machine Learning Classifiers

Although Deep Learning text classifiers show very strong and stable results on imbalanced classification tasks as compared to linear classifier (*37*), our first experiments with an early training data set based on PLOS articles resulted in f-score of 96.3% for mentioning citations, 55.3% for supporting, and 20.5% for disputing. The initial accuracy for disputing in particular raised concerns about the feasibility of the task itself at scale. We focused on multiple approaches to increase over time the accuracy of classifier for the two less frequent classes:

‐ Improving the classification architecture: After initial experiments with RNN architectures (BidGRU), we obtained significant improvements with ELMo dynamic embeddings (*38*) and an ensemble approach. Although the first experiments with BERT (*39*) were disappointing, fine-tuning SciBERT (*40*) led to the best results and is the current production architecture of the platform.
‐ Using oversampling and class weighting techniques: It is known that the techniques developed to address imbalanced classification in traditional ML can be applied successfully to Deep Learning too (*41*). We introduced in our system oversampling of less frequent classes, class weighting, and meta-classification with three binary classifiers. These techniques provide some improvements, but they rely on empirical parameters which must be re-evaluated as the training data changes.
‐ Extending the training data for less-frequent classes: As mentioned previously, we use an active learning approach to select the likely less frequent citation classes based on the scores of the existing classifiers. By focusing on edge cases over months of manual annotations, we observed significant improvements in performance for predicting disputing and supporting cases.

Table 1 presents the model evaluation after iterations of the classification system over time using our fixed holdout set. Table 2 presents the evaluation metrics for the current SciBERT model. Reported scores are averaged over 10 runs. The F-score for the classification of “disputing” was notably improved from 20.1% to 58.97%. The precision of prediction of “disputing” in particular reaches 85.19%, a very reliable level for such a rare class.

**Table 1.**
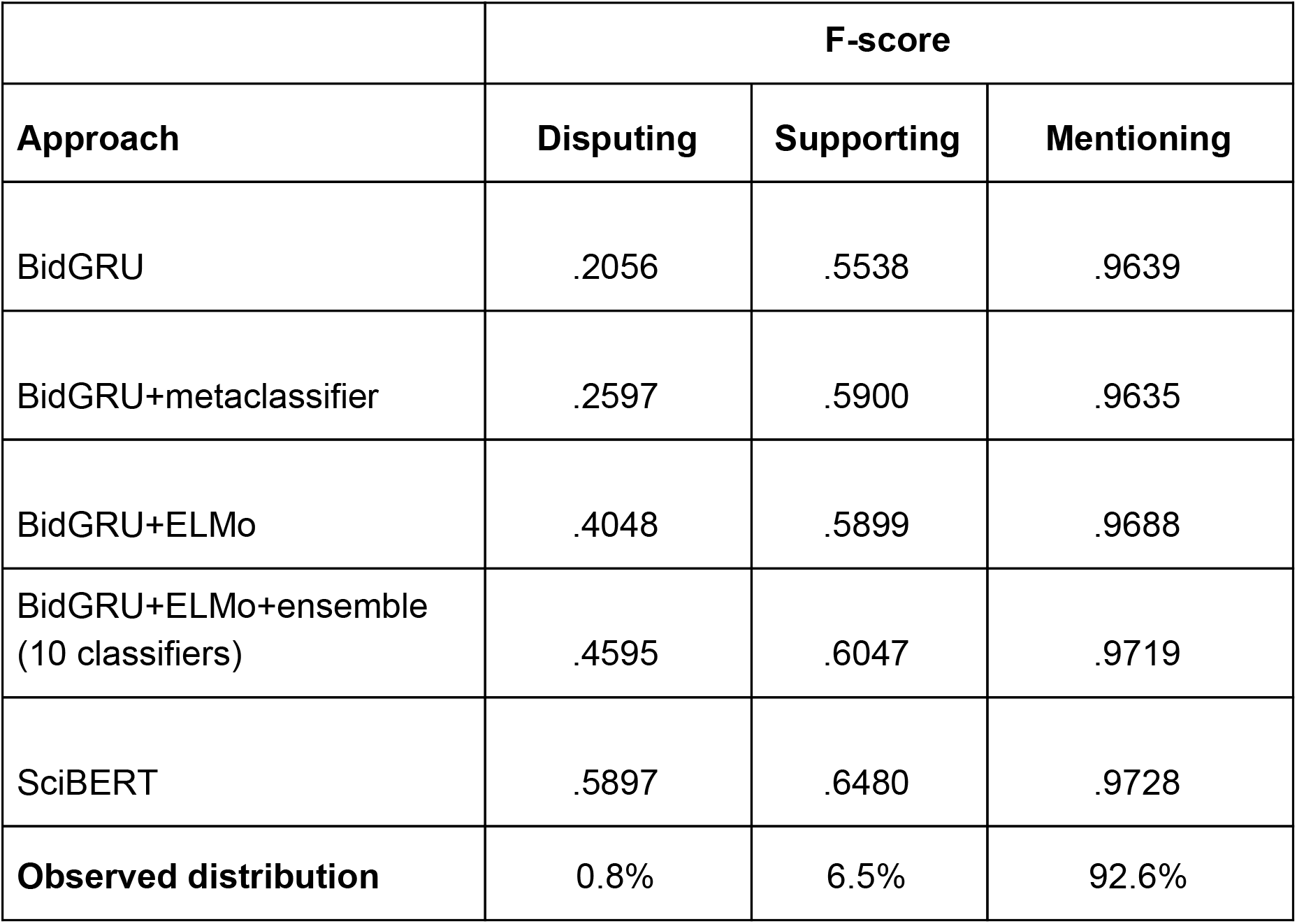
Progress on classification results over approx. 1 year, evaluated on a fixed holdout set of 9,708 examples. In parallel to these various iterations on the classification algorithms, the training data was raised from 30,665 (initial evaluation with BidGRU) to 38,925 examples (last evaluation with SciBERT) via an active learning approach.

|  | F-score |  |  |
| --- | --- | --- | --- |
| Approach | Disputing | Supporting | Mentioning |
| BidGRU | .2056 | .5538 | .9639 |
| BidGRU+metaclassifier | .2597 | .5900 | .9635 |
| BidGRU+ELMo | .4048 | .5899 | .9688 |
| BidGRU+ELMo+ensemble<br>(10 classifiers) | .4595 | .6047 | .9719 |
| SciBERT | .5897 | .6480 | .9728 |
| **Observed distribution** | 0.8% | 6.5% | 92.6% |

**Table 2.** Accuracy of SciBERT classifier, currently deployed on the scite platform, evaluated on a holdout set of 9,708 examples. Note: when deploying classification models in production, we balance the precision/recall so that all the classes have a precision higher than 80%.

|  | **Precision** | **Recall** | **F-score** |
| --- | --- | --- | --- |
| Disputing | .8519 | .4510 | .5897 |
| Supporting | .7406 | .5760 | .6480 |
| Mentioning | .9615 | .9843 | .9728 |

Given the unique nature of scite, there are a number of additional considerations. First, scaling is a key requirement of scite, which addresses the full corpus of scientific literature. While providing good results, the prediction with the ELMo approach is 20 times slower than with SciBERT, making it less attractive for our platform. Second, we have experimented with using section titles to improve classifications – for example, one might expect to find supporting and disputing statements more often in the Results section of a paper, and mentioning statements in the Introduction. Counterintuitively, including this information in our model had no impact on F-scores, although it did slightly improve precision. Third, segmenting scientific text into sentences presents unique challenges due to the prevalence of abbreviations, nomenclatures, and mathematical equations. Finally, we experimented with various context windows (i.e., the amount of text used in the classification of a citation), but were only able to improve the F-score for the disputing category by 8 points by manually selecting the most relevant phrases in the context window. Automating this process might improve classifications, but doing so presents a significant technical challenge. Other perspectives of improvement of the classifier include multitask training, refinement of classes, increase of training data via improved active learning techniques, and integration of categorical features in the transformer classifier architecture.

We believe that the specificity of our evidence-based citation classes, the size and the focus on the quality of our manually annotated dataset (multiple rounds of blind-annotations with final collective reconciliation), the customization and continuous improvement of a state of the art Deep Learning classifier, and finally the scale of our citation analysis distinguishes our work from existing developments in automatic citation analysis.

### Citation statement and classification pipeline

TEI XML data is parsed in Python using the BeautifulSoup library and further segmented into sentences using a combination of Spacy (*42*) and Natural Language Toolkit’s Punkt Sentence Tokenizer (*43*). These sentence segmentation candidates are then post-processed with custom rules to better fit scientific texts, existing text structures and inline markups. For instance, a sentence split is forbidden inside a reference callout, around common abbreviations not supported by the general-purpose sentence segmenters, or if it is conflicting with a list item, paragraph, or section break.

The implementation of the classifier is realized by a component called *Veracity*, providing a custom set of deep learning classifiers built on top of the Open Source DeLFT library (*44*). Veracity is written in Python and employs Keras and TensorFlow for text classification. It runs on a single server with an NVIDIA GP102 (GeForce GTX 1080 Ti) graphics card with 3584 CUDA cores. This single machine is capable of classifying all citation statements as they are processed. Veracity retrieves batches of text from the scite database that have yet to be classified, processes them, and updates the database with the results. When deploying classification models in production, we balance the precision/recall so that all the classes have a precision higher than 80%. For this purpose, we use the holdout dataset to adjust the class weights at the prediction level. After evaluation, we can exploit all available labeled data to maximize the quality, and the holdout set captures a real-world distribution adapted to this final tuning.

### User Interface

The resulting classified citations are stored and made available on the scite platform. Data from scite can be accessed in a number of ways (downloads of citations to a particular paper; the scite API, etc.). However, users will most commonly access scite through its web interface. Scite provides a number of core features, detailed below.

The scite report page (Figure 2) displays summary information about a given paper. All citations in the scite database to the paper are displayed, and users can filter results by classification (supporting, mentioning, disputing), paper section (e.g., Introduction, Results), and the type of citing article (e.g., preprint, book, etc.). Users can also search for text within citation statements and surrounding citation context. For example, if a user wishes to examine how an article has been cited with respect to a given concept (e.g., fear), they can search for citation contexts that contain that key term. Each citation statement is accompanied by a classification label, as well as an indication of how confident the model is of said classification. For example, a citation statement may be classified as supporting with 90% confidence, meaning that the model is 90% certain that the statement supports the target citation. Finally, each citation statement can be flagged by individual users as incorrect, so that users can report a classification as incorrect, as well as justify their objection. After a citation statement has been flagged as incorrect, it will be reviewed and verified by two independent reviewers, and, if both agree, the recommended change will be implemented. In this way, scite supplements machine learning with human interventions to ensure citations are accurately classified.

In order to improve the utility and usability of the smart citation data, scite offers a wide variety of tools common to other citation platforms such as Scopus and Web of Science and other information retrieval software. These include literature searching functionality for researchers to find supported and disputed research, visualizations to see research in context, reference checking for automatically evaluating references with scites data on an uploaded manuscript and more. scite also offers plugins for popular web browsers and reference management software (e.g., Zotero) that allow easy access to scite reports and data in native research environments. Finally, scite provides citation indices that rank and evaluate journals and funders based on aggregate smart citation data in order to provide alternatives to the journal impact factor during research evaluation.

## Discussion

### Research Applications

A number of researchers have already made use of scite for quantitative assessments of the literature. For example, Bordignon (*45*) examined self-correction in the scientific record and operationalized “negative” citations as those which scite classified as disputing. They found that negative citations are rare, even among works that have been retracted. In another example, Nicholson et al (*46*) examined scientific papers cited in Wikipedia articles and found that – like the scientific literature as a whole – the vast majority presented findings that have not been subsequently verified. Similar analyses could also be applied to articles in the popular press.

One can imagine a number of additional, metascientific applications. For example, network analyses with directed graphs, valenced edges (by type of citation – supporting, disputing, and mentioning), and individual papers as nodes could aid in understanding how various fields and subfields are related. A simplified form of this analysis is already implemented on the scite website (see Figure 4), but more complicated analyses that assess traditional network indices such as centrality, clustering, etc. could be easily implemented using standard software libraries and exports of data using the scite API.

**Figure 4.**
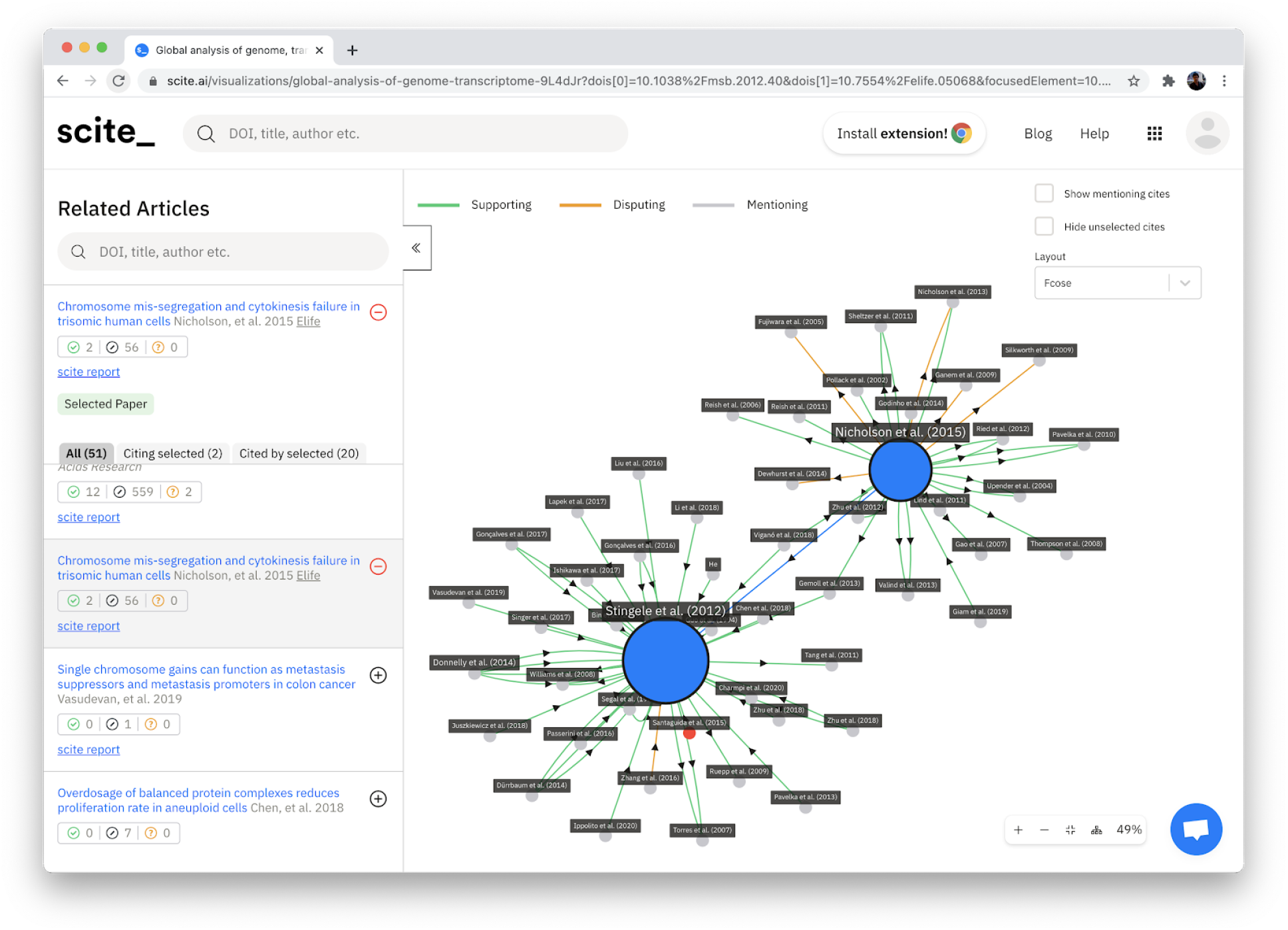
A citation network representation using the scite Visualization tool. The nodes represent individual papers, with the edges representing supporting (green) or disputing (orange) citation statements. The graph is interactive and can be expanded and modified for other layouts.

### Implications for scholarly publishers

There are a number of implications for scholarly publishers. At a very basic level, this is evident in the features scite provides that are of particular use to publishers. For example, the scite Reference Check parses the reference list of an uploaded document and produces a report indicating how items in the list have been cited, and flagging those which have been retracted or have otherwise been the subject of editorial concern. This type of screening can help publishers and editors ensure that articles appearing in their journals do not inadvertently cite discredited works. Evidence in scite’s own database indicates that this would solve a seemingly significant problem, as in 2019 alone, nearly 6,000 published papers cited works that had been retracted prior to 2019. Given that over 95% of citations made to retracted articles are in error (*47*), had the Reference Check tool been applied to these papers during the review process, the mistakes could have been caught.

However, there are additional implications for scholarly publishing that go beyond the features provided by scite. We believe that by providing insights into how articles are cited - rather than simply noting that the citation has occurred – scite can alter the way journals, institutions, and publishers are assessed. Scite provides journals and institutions with dashboards that indicate the extent to which papers with which they are associated have been supported or disputed by subsequent research. Even without reliance on specific metrics, the approach scite provides begs the question: what if we normalized the assessment of journals, institutions and researchers in terms of how they were cited rather than the simple fact that they were cited alone?

### Implications for researchers

Given the fact that nearly 3 million scientific papers are published every year (*48*), researchers increasingly report feeling overwhelmed by the amount of literature they must sift through as part of their regular workflow (*49*). Scite can help by assisting researchers in identifying relevant, reliable work that is narrowly tailored to their interests, as well as to better understand how a given paper fits into the broader context of the scientific literature. For example, one common technique for orienting oneself to new literature is to seek out the most highly cited papers in that area. However, if the context of those citations are also visible, the value of a given paper can be more completely assessed. In other words, scite allows its users to not only search through references provided by a given paper but also review the true scientific impact this paper has made.

There are, however, additional – although perhaps less obvious – implications. If citations are easily visible, it is possible that researchers will be incentivized to make replication attempts easier (for example, by providing more explicit descriptions of methods, instruments, etc.) in hopes that their work will be replicated.

### Limitations

At present, the biggest limitation for researchers using scite is the size of the database. At the time of this writing, scite has ingested over 800 million separate citation statements from over 23 million scholarly publications. However, there are over 70 million scientific publications in existence (*48*). scite is constantly ingesting new papers from established sources and signing new licensing agreements with publishers, so this limitation should abate over time. However, given that the ingestion pipeline fails to identify approximately 30% of citation statements/references in PDF files (∼5% in XML), the platform will necessarily contain fewer references than services like Google Scholar and Web of Science, which do not rely on ingesting the full text of papers. Even if references are reliably extracted and matched with a DOI or directly provided by publishers, a reference is currently only visible on the scite platform if it is matched with at least one citation context in the body of the article. As such, the data provided by scite will necessarily miss a measurable percentage of citations to a given paper, although this number will decrease over time as more full-text XML are available and PDF document structuring techniques are improving.

Another limitation is related to the classification of citations. First, as noted previously, the Veracity software does not perfectly classify citations. The accuracy of the classifier will likely increase over time as technology improves and the training dataset increases in size. Second, the ontology currently employed by scite (supporting, mentioning, and disputing) necessarily misses some nuance regarding how references are cited in scientific papers. One key example relates to what “counts” as a disputing citation: at present, this category is limited to instances where new evidence is presented (e.g., a failed replication attempt or a difference in findings). However, it might also be appropriate to include conceptual and logical arguments against a given paper in this category.

## Conclusions

The automated extraction and analysis of scientific citations is a technically challenging task, but one whose time has come. By surfacing the context of citations rather than relying on their mere existence as an indication of a paper’s importance and impact, scite provides a novel approach to addressing pressing questions for the scientific community, including incentivizing replicable works, assessing an increasingly large body of literature, and quantitatively studying entire scientific fields.

## Acknowledgments

We would like to thank Yuri Lazebnik for his help in conceptualizing and building scite.

## Author Contributions

JMN was involved in conception and design, acquisition of data, analysis and interpretation of data, and drafting or revising the article. MM was involved in acquisition of data and analysis and interpretation of data. PL was involved in conception and design, analysis and interpretation of data and and drafting or revising the article. AU was involved in analysis and interpretation of data and drafting or revising the article. DR was involved in analysis and interpretation of data and drafting or revising the article. NPR was involved in conception and design. SCR was involved in conception and design, acquisition of data, analysis and interpretation of data, and drafting or revising the article.

## Data availability

Code used in the ingestion of manuscripts is available at https://github.com/kermitt2/grobid and https://github.com/kermitt2/biblio-glutton. The classification of citation statements is performed by a modified version of DeLFT (https://github.com/kermitt2/delft). The training data used by the scite classifier is proprietary and not publicly available. The 800+ million citation statements are available at scite.ai but can not be shared in full due to licensing arrangements made with publishers and copyright restrictions.

## Footnotes

1 As an illustration, the ISTEX project has been an effort from the French state leading to the purchase of 23 million full text articles from the mainstream publishers (Elsevier, Springer-Nature, Wiley, etc.) mainly published before 2005, corresponding to an investment of €55 million in acquisitions. The delivery of full text XML when available was a contractual requirement, but an XML format with structured body could be delivered by publishers for only around 10% of the publications.

2 For more information on the history and prevalence of Crossref, see https://www.crossref.org/about/

